# Human Foot Force Informs Balance Control Strategies when Standing on a Narrow Beam

**DOI:** 10.1101/2024.02.27.582256

**Authors:** Kaymie Shiozawa, Marta Russo, Jongwoo Lee, Neville Hogan, Dagmar Sternad

## Abstract

Despite the abundance of studies on the control of standing balance, insights about the roles of biomechanics and neural control have been limited. Previous work introduced an analysis combining the direction and orientation of ground reaction forces. The “intersection point” of the lines of actions of these forces exhibited a consistent pattern across healthy, young subjects when computed for different frequency components of the center of pressure signal. To investigate the control strategy of quiet stance, we applied this intersection point analysis to experimental data of 15 healthy, young subjects balancing in tandem stance on a narrow beam and on the ground. Data from the sagittal and frontal planes were analyzed separately. The task was modeled as a double-inverted pendulum controlled by an optimal controller with torque-actuated ankle and hip joints and additive white noise. To test our prediction that the controller that minimized overall joint effort would yield the best fit across the tested conditions and planes of analyses, experimental results were compared to simulation outcomes. The controller that minimized overall effort produced the best fit in both balance conditions and planes of analyses. For some conditions, the relative penalty on the hip and ankle joints varied in a way relevant to the balance condition or to the plane of analysis. These results suggest that unimpaired quiet balance in a challenging environment can be best described by a controller that maintains minimal effort through the adjustment of relative ankle and hip joint torques.

**NEW & NOTEWORTHY:** This study explored balance control in humans during a challenging task using the novel intersection point analysis, based on ground reaction force direction and point of application. Experimental data of subjects standing on a narrow beam in tandem stance were compared with modeling results of a double-inverted pendulum. The analysis showed that individuals minimized effort by adjusting ankle and hip torques, shedding light on the interplay of biomechanics and neural control in maintaining balance.

## INTRODUCTION

Despite the abundance of studies on postural balance, the control mechanisms of this complex whole-body task are still being discussed (1–3). Analyzing the dynamics of quiet standing has proven limited, as the only available ‘input’ is sensorimotor noise that is internal to the biological system and cannot be directly measured (4–7). To overcome this challenge, numerous studies have applied external mechanical perturbations to standing humans as input for the identification of balance control mechanisms (2, 8, 9). Nevertheless, quiet standing experiments allow for observations of natural stance and longer-term recordings. One form of mild and safe system interrogation is to expose human subjects to a challenging environment without applying perturbations (10–13). The present study examined human subjects standing on a narrow beam in tandem stance to gain insights into the control mechanisms of postural balance.

To conduct closed-loop system identification of quiet stance, the output should contain sufficient information about the dynamics of the system. Prior work studying quiet or perturbed balance has typically used the center of pressure, the point of application of the ground reaction force, as a way to quantify subjects’ balance ability (6, 14–17). However, the available force data contain more information than its point of application. The orientation of the ground reaction force enriches the information for our analyses, because the horizontal component of the ground reaction force is related to the lateral acceleration of the center of mass (18). Even though this shear component is dwarfed by the vertical component, the horizontal force is essential to provide translational stability. With this additional measurement, the lines of action of the ground reaction forces can be computed. The “intersection point” of these forces’ lines of action can provide informative details about the dynamics and control of human balance (18).

First measured in quiet standing by Gruben and colleagues (18), the intersection point was defined as the point where the lines of action of foot-ground forces in a plane exhibited minimal deviation. The height of this intersection point can shed light on the contributions of biomechanics and neural control. When the intersection point is above the center of mass, the body behaves like a single rigid body, i.e., stiffening the hip and rotating about the ankle. In contrast, when the intersection point is below the center of mass or at the ground, ankle torque is used less or not at all, as the body is only stabilized by hip movements. Previous work showed that when the intersection point height was calculated for different frequency bands of the center of pressure, there was a consistent pattern in healthy, young subjects in quiet stance (18).

Following these first insights, a recent study by Shiozawa and colleagues (19) aimed to account for the intersection point’s frequency-dependent pattern with a control model. The sagittal plane data of humans quietly standing on the ground were compared to numerical simulation results based on a mathematical model of a double-inverted pendulum controlled by a well-established optimal control model. The results suggested that subjects’ control strategy was best described by a controller that minimized the control effort at the ankle and hip joints, with greater penalty on using hip torque.

The current study aimed to determine whether the principle of minimum effort also governed control in challenging balance conditions, i.e., tandem stance on a narrow beam. This condition was compared against the same stance on the ground. Furthermore, because on the beam the mediolateral direction requires more stabilization than the anteroposterior direction (20), this work also analyzed the intersection point in both the sagittal and frontal planes. We were additionally interested in how the relative roles of the ankle and hip would differ across the conditions and across the planes of analyses in the context of minimized effort.

Consistent with previous results (19), we predicted that the controller that minimized overall joint effort would yield the best fit across the tested conditions and planes of analyses. To test this prediction, we tasked 15 subjects to balance in tandem stance on a narrow beam and on the ground. The experimental data were processed using the same analysis as before to examine the frequency-dependent intersection points for each balance condition. Assuming that a quietly standing human can be adequately modeled using a double-inverted pendulum with torque-actuated ankle and hip joints corrupted by white noise, an optimal control method was employed to simulate a human standing upright. Best-fit controller parameters that quantified unimpaired human control strategies under challenged balance conditions were identified by comparison with the experimental results.

Results of the simulation showed that, similar to previous outcomes, the minimum-effort controller described the human data well across all tested conditions and planes of analyses. When balancing on the beam, ankle torque in the frontal plane was penalized more than hip torque. The opposite behavior was observed when subjects stood on the ground. The relative penalty on the joint torques in the sagittal plane was skewed towards the hip. These findings underscored our simple model’s ability to address a range of stance conditions under a single overarching principle: the economization of effort.

## MATERIALS AND METHODS

### Human Experiment

#### Participants

Fifteen participants (eight males and seven females, between 19 and 36 years) with no history of neurological conditions took part in the experiment. The subjects’ average mass and height were 69 ± 16 kg and 1.66 ± 0.10 m, respectively. The study was approved by the Institutional Review Board of Northeastern University in accordance with the Declaration of Helsinki (IRB# 18-01-19).

#### Apparatus and Setup

Participants stood on a narrow wooden beam (width 3.65 cm, height 7.62 cm) that was placed on top of a six-axis force plate (AMTI, Watertown, MA, USA), as shown in Figure 1. For comparison, subjects also stood on the ground directly on the force plate. The ground reaction force data were measured at 500 Hz. Further details of the experimental apparatus and setup can be found in the previous study by Russo et al. (10).

**Figure 1.**
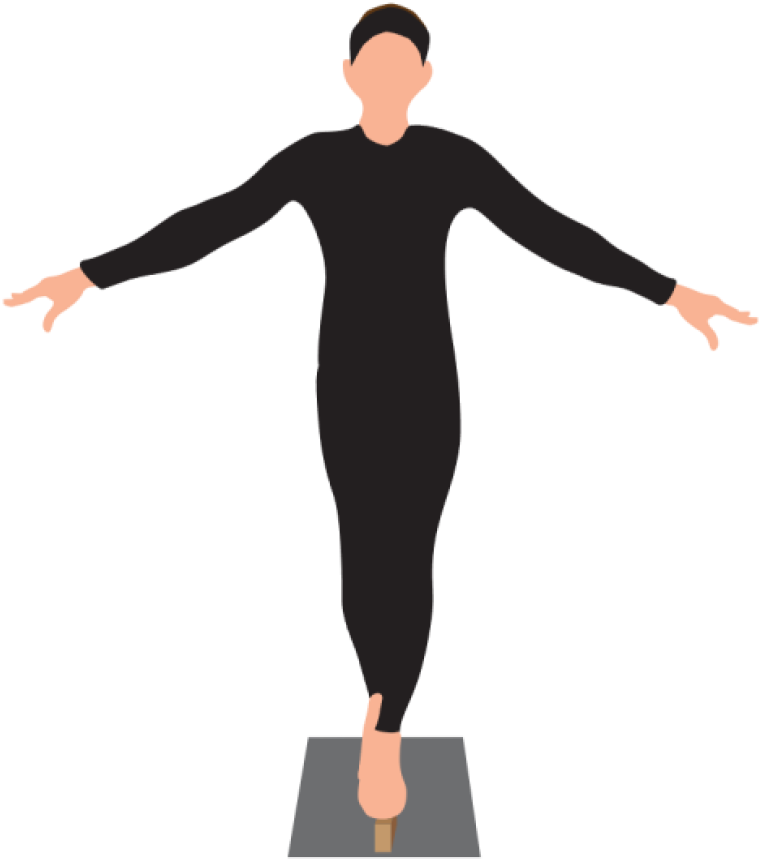
Experimental setup. For each balance condition, subjects were tasked to stand in tandem stance with their arms extended out to the side. Subjects either stood on a narrow beam or on the ground. A force plate was placed under the beam or directly under the subjects’ feet depending on the condition.

#### Protocol

Participants were asked to stand barefoot in tandem stance while gazing straight ahead viewing a point marked on the wall (5 m away, Figure 1). Subjects were free to decide which foot to place in front of their stance, but then maintained that position for all trials. They were also allowed to freely move their arms to maintain balance. Two platform conditions, one on the beam and the other on the ground, were tested. Subjects performed three trials on the beam and two trials on the ground; each trial lasted 30 s. For the analysis, the first and last three seconds were deleted to exclude transient effects; thus, each analyzed trial had 24 s worth of data. Subjects could step off the beam between the trials. However, if the subjects stepped out of tandem stance during a trial, that trial was discarded. The subjects were not permitted to redo the trial, as repeatedly performing the task could induce fatigue. Overall, out of 75 trials, 13 trials were discarded (17%).

#### Intersection Point Analysis

Previous work defined the intersection point as a spatially fixed point in a plane where the net ground reaction force vectors minimally deviated from each other (18), as depicted in Figure 2a. The present study adopted the same assumptions and mathematical definitions, which are briefly outlined.

**Figure 2.**
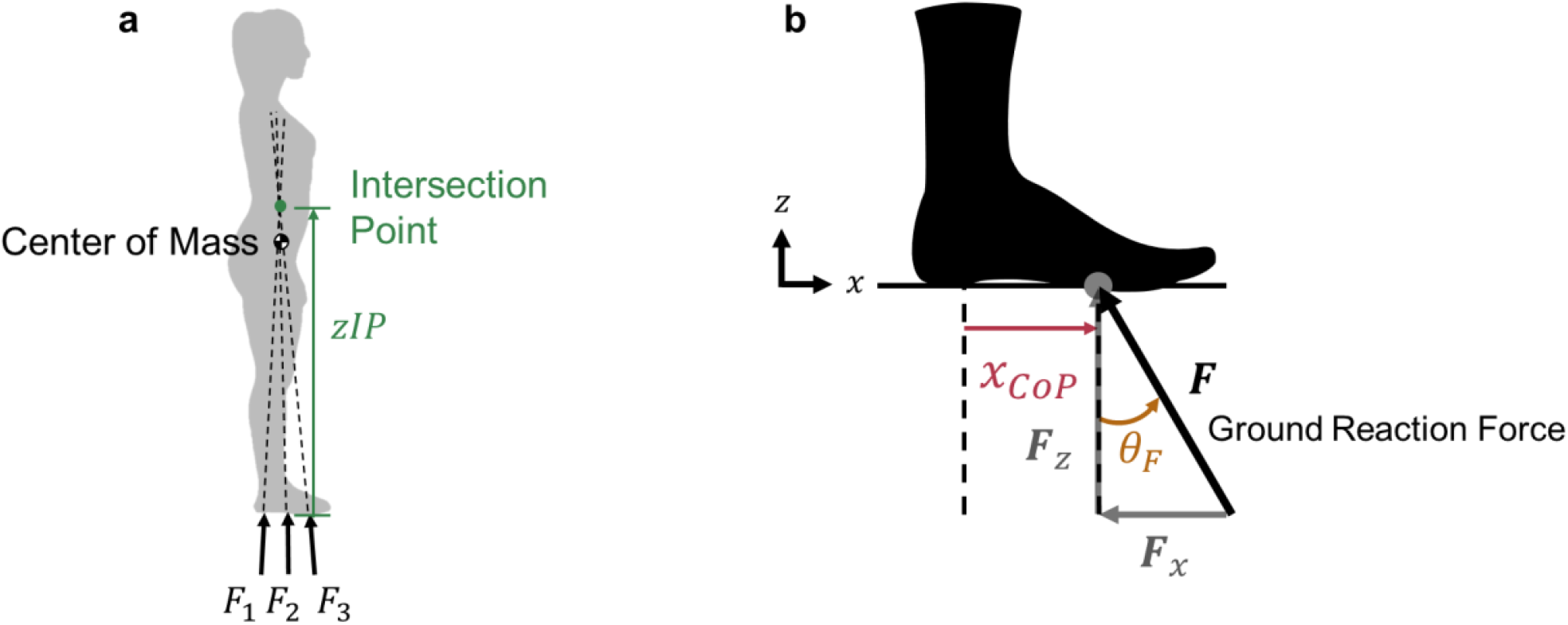
a) The intersection point was defined in previous work as a spatially fixed point in a plane where the net ground reaction force vectors (e.g. *F* _1_, *F* _2_, *F* _3_) minimally deviated from each other (18) (figure adapted from Boehm et al.). The height of the intersection point (*z*_*IP*_) and the center of mass location is indicated as well. **b)** Illustration of the center of pressure position (*x* _*CoP*_) and force angle (*θ* _*F*_) signals with respect to the ground reaction force vector (***F***). Vertical (***F***_*z*_) and horizontal (***F*** _x_) force components are shown as well. The center of pressure position’s reference at any time was the reference point of the force plate.

It was assumed that body movements and variations in the ground reaction force directions were both small to allow the foot-force orientation (*θ* _*F*_) to be approximated as

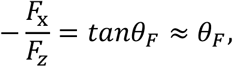

where *F*_x_ and *F*_*z*_ are the horizontal and vertical components of the ground reaction force vector (***F***). Figure 2b depicts these two signals.

To calculate the intersection point height, both *θ* _*F*_ and *x* _*CoP*_ were first analyzed in the frequency domain to investigate the system response. To conduct this frequency-domain analysis, a Hamming window with the length of the data was applied to both the *θ* _*F*_ and *x* _*CoP*_ signals. These signals were then bandpass-filtered (zero-lag, 2^nd^-order Butterworth) into bands of 0.2 Hz width, centered on frequencies from 0.5 to 7.9 Hz (38 nominally non-overlapping bands). The height of the intersection point at each frequency band was the slope of the principal eigenvector of the best-fit covariance matrix of the filtered *x* _*CoP*_ plotted against filtered *θ* _*F*_, as shown in Figure 3. The intersection point height (*zIP*) could be expressed as

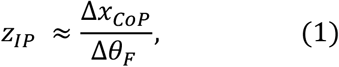

**Figure 3.**
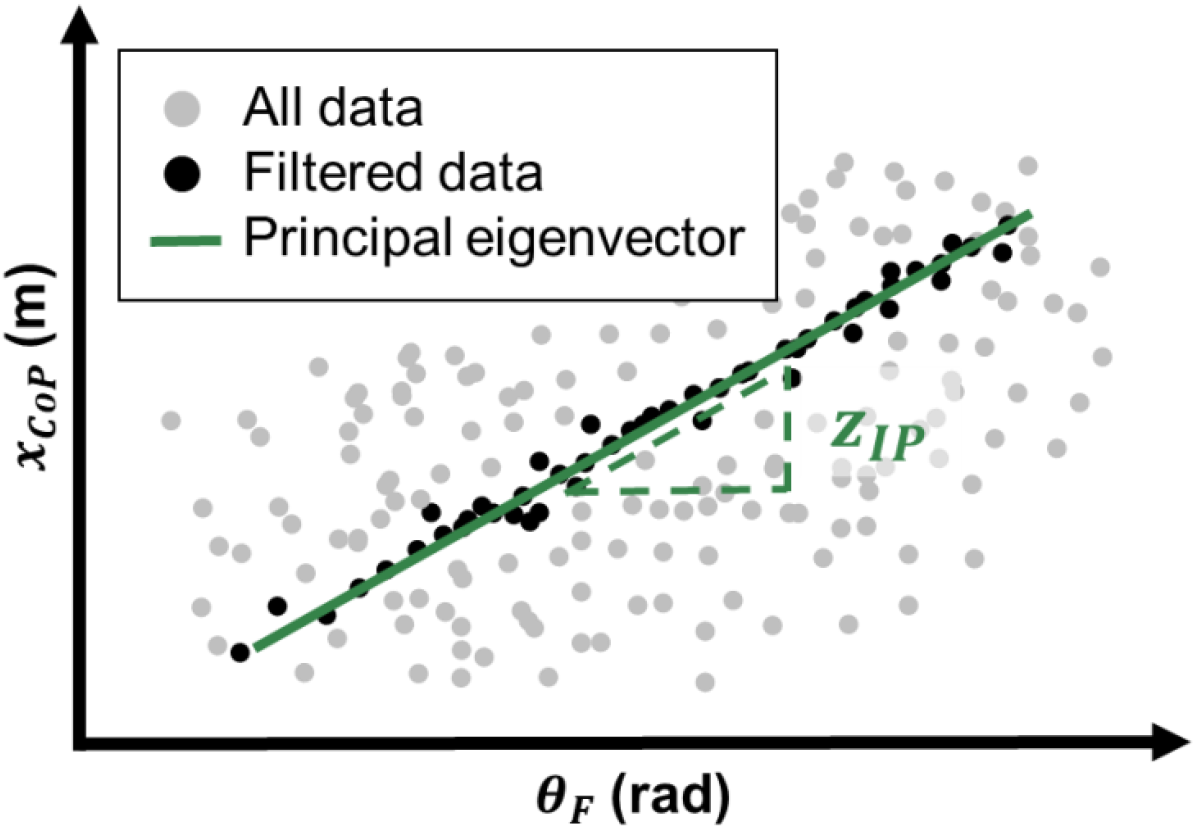
Graphical illustration of how the intersection point height (*z* _*IP*_) was determined by the slope of the principal eigenvector of the relation between filtered *x* _*CoP*_ and *θ* _*F*_. The gray points represent all data points of a single trial’s time series, the black points represent the bandpass-filtered data at a given frequency band, and the green line represents the principal eigenvector of the filtered data. The slope of the principal eigenvector corresponds to *z* _*IP*_, as indicated by the relationship from Equation 1.

where Δ*x* _*CoP*_ is a small change in the filtered center of position and *θ* _*F*_ is a small change in filtered ground reaction force orientation. The intersection point was obtained separately in the sagittal and frontal planes, using the mediolateral and anteroposterior components of the *x* _*CoP*_ .

For each balance condition, the processed *z*_*IP*_ data were plotted against their respective frequency band. *z*_*IP*_ was normalized by the center of mass height per subject and the means at each frequency band were calculated. Values of the *z*_*IP*_ that were three standard deviations away from the overall mean were discarded (< 5% of the data).

### Simulation

#### Model

Previous work (19) showed that a single inverted pendulum model, though still widely used in balance studies (9, 21, 22), was unable to replicate the intersection point below the center of mass that was observed in humans. Therefore, the next simplest model, a double-inverted pendulum model, was employed in this study. To represent the T-shaped posture (arms extended to the sides) with which subjects balanced, this model attached rods orthogonal to the upper pendulum to emulate the extended arms. For this study, the arms remained fixed at the orthogonal angle (even though some subjects moved their arms). The double-inverted pendulum model that was used to simulate a multi-segmented human body in the sagittal and frontal planes is illustrated in Figure 4.

**Figure 4.**
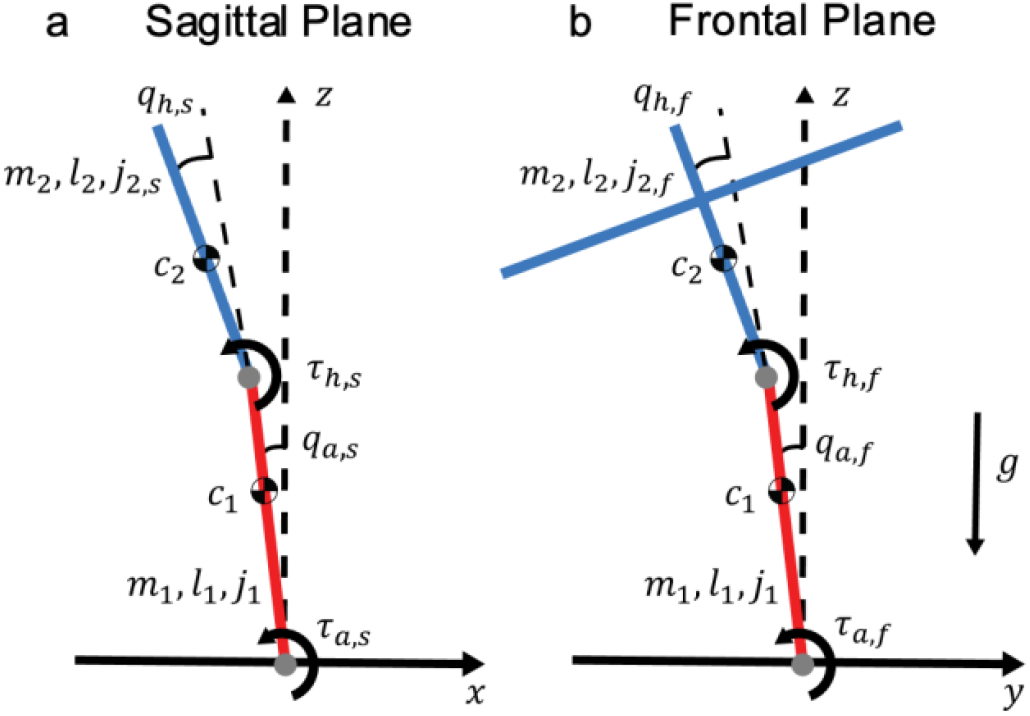
Double inverted pendulum model in **a)** the sagittal plane and **b)** the frontal plane. The bottom red segment represents the lower body and the blue link above it reflects the upper body including the rigidly extended arms. The labels indicate angle (*q* _*a*_, ankle; *q* _*h*_, hip) and torque (*τ* _*a*_, ankle; *τ* _*h*_, hip) conventions and parameter values for mass (*m*_*i*_), length (*l*_*i*_), center of mass position (*c*_*i*_), and moment of inertia about the center of mass (*j*_*i*_). Subscripts *s* and *f* indicate the sagittal and frontal planes’ parameters respectively, but they have been omitted in the main body of this paper for brevity. The direction of gravity (*g*) is also illustrated.

Based on the anthropometric distribution of male subjects (23), the lumped model parameters were determined (Table 1). Preliminary analysis showed that the results did not differ when the anthropometric distribution of female subjects was used. Any mass and length below the ankle were neglected, as the model assumed the ankle to be a pin joint. The center of mass positions were measured with respect to the ankle joint for the lower body (link 1) and with respect to the hip joint for the upper body (link 2). The moments of inertia were calculated about the center of mass of each link. The equations of motion, the orientation of the ground reaction force vector, and the center of pressure position, which were all required to quantify the *z*_*IP*_, were calculated in a similar manner as in the previous study (19). The lumped parameters were replaced by those shown in Table 1. The states and input variables were **x = [*q, dq***]^*T*^, ***q* = [***q*_*a*_, *q*_*h*_]^*T*^, and ***τ* = [***τ* _*a*_, *τ*_*h*_]^*T*^ as defined in Figure 4.

**Table 1.** Lumped model parameters.

| Symbol | Parameter (units) | Plane | Value |  |
| --- | --- | --- | --- | --- |
|  |  |  | Link 1<br>Lower Body | Link 2<br>Upper Body |
| $m$ | Mass (kg) | | 25.33 | 41.29 |
| $l$ | Length (m) | | 0.822 | 0.807 |
| $l_c$ | Center of mass (m) | | 0.559 | 0.362 |
| $j$ | Moment of inertia (kgm<sup>2</sup>) | Sagittal | 1.213 | 1.809 |
|  |  | Frontal |  | 3.587 |
| $g$ | Gravitational acceleration (m/s<sup>2</sup>) | | 9.81 | |

The noise processes that were added to the ankle and hip joint torques were assumed to be white, mutually uncorrelated, and followed a zero-mean Gaussian distribution, with standard deviations for the ankle and hip denoted as 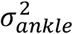 and 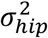, respectively. The relative strength of the two noise sources was defined as 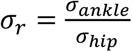.

#### Controller

To guarantee a stable closed-loop system, the linear quadratic regulator (LQR) was used. The double-inverted pendulum model was nonlinear, but the controller was linear; therefore, the nonlinear equations of motion were first linearized about the upright posture: **x**_0_ **= [**0, 0, 0, 0]^*T*^ and ***τ*** _0_ **= [**0, 0]^*T*^. The LQR computes an optimal control gain matrix based on the quadratic cost function

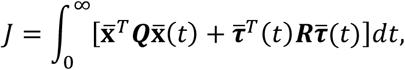

where 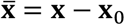 and 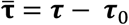 and ***Q*** and ***R*** weight the state and input deviations from zero. The input weighting matrix was parameterized as

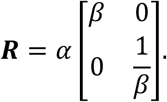

When *α* was large, the control effort was reduced to a minimum value required for stability. On the other hand, when *β* > 1, the ankle torque was penalized more heavily than the hip, and vice versa when *β* < 1.

The LQR approach guaranteed stability for all values of *α* and *β*. This fact justified the use of a linear method to study a nonlinear system. Previously, the effect of *α* on the intersection point was explicitly analyzed (19), and a large *α* was shown to best fit healthy human subject data. To further test the hypothesis that unimpaired balance was best described by a controller that minimized effort, *α* was set to be either very large (10^6^) or very small (10^−4^). With large *α*, the closed loop system exhibited a predetermined behavior by situating its poles at the mirrored positions (across the imaginary axis) of the unstable open-loop poles; this behavior was unaffected by the specific choice of the state weighting matrix ***Q***, as long as ***R*** was sufficiently larger than ***Q***. Although the poles did not vary significantly for large *α*, the closed loop gain matrix varied considerably with *β*.

#### Simulation Protocol

To quantify the variation of best-fit parameters under different tested balance conditions, parameters were varied using the following procedure. First, the parameter *α* that weighted the relative cost of the control input was set to a large (10^6^) value. Then, a global parameter search of the parameter that adjusted the relative cost on the hip and ankle torques, *β*, and the joint noise ratio, *σ* _*r*_, was conducted for that *α*. For *β*, values between 0.01 and 3 were explored; for *σ* _*r*_, values between 0.01 and 20 were explored. These ranges were selected based on a preliminary analysis that evaluated the numerical stability of the simulation at different *β* and *σ* _*r*_ values. The search procedure was repeated for small *α* (10^−4^). To search for the best fit *α, β*, and *σ* _*r*_ parameters, an analytic approach was used to compute the intersection point height and facilitate faster iteration through the parameters (24).

Once the best-fit parameters were determined, 40 simulation trials were conducted for each balance condition to enable statistical analysis of the simulated dependence of the intersection point height on frequency. The simulation was conducted using semi-implicit Euler integration. The initial condition was set to **x**0. Replicating the experimental protocol, each simulation was run for 24 s with a time step of 2 ms (500 Hz). Because human balance has only small vertical motions, the height of the center of mass was calculated based on the subjects’ average height. This constant center of mass height was used to normalize the intersection point height for each subject. All simulations were conducted in MATLAB 2022b (Mathworks, Natick MA).

For each simulation trial, the control input at each time step was stored to compute the root-mean-squared (RMS) value of the torque for the ankle and hip joints separately. Then, the ratio of the joints’ RMS torques was computed as

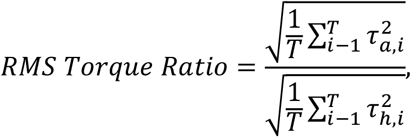

where *τ* _*a*_ and *τ* _*h*_ are ankle and hip torques, and *T* is the number of time steps. Finally, across the 40 simulation runs for the best-fit *β* and *σ* _*r*_ parameters, the mean and standard deviation of the RMS torque ratio were reported.

#### Comparison of Experimental and Simulated Results

To determine the best fit across different model parameters, the root-mean-squared errors (RMSE) between the computed and experimental data for each balance condition were calculated across all frequency bands by

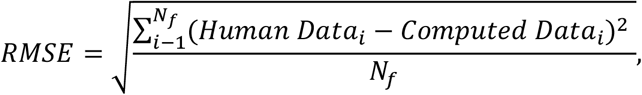

where *Human Data* _*i*_ was the average observed intersection point height normalized by each individual’s center of mass height for each balance condition in a given frequency band. For the global parameter search, *Computed Datai* was the analytically computed intersection point height (24) normalized by a constant center of mass height in a given frequency band. *N* _*f*_ was set to be 38 for this study, as there were 38 different frequency bands in which the intersection point height was computed. The parameter set with the lowest *RMSE* value across all tested *α, β*, and *σ* _*r*_ was selected as the best-fit parameter set for that balance condition. Once the best-fit parameters were determined, the *RMSE* for each balance condition was recomputed by setting *Computed Datai* as the average intersection point height normalized by a constant center of mass height across 40 simulation trials in a given frequency band.

## RESULTS

### Human Results

Figure 6 shows the mean heights of the intersection point for the beam and ground condition over different frequency bands; the data are shown separately for the sagittal and frontal planes. At first glance, the intersection point varies with frequency in a similar manner when subjects stood on the beam and on the ground in both planes: the intersection point was above the center of mass at low frequencies and below the center of mass at high frequencies. In particular, very similar patterns for standing on or off the beam emerged in the sagittal plane, even though the variability was smaller at lower frequencies when standing on the ground (Figure 6a and 6b). The intersection point heights were close to the center of mass at similar frequencies: at 1.5 Hz (beam) and 1.3 Hz (ground). Furthermore, the mean of the intersection point height at all frequencies was always above 0.45.

**Figure 6.**
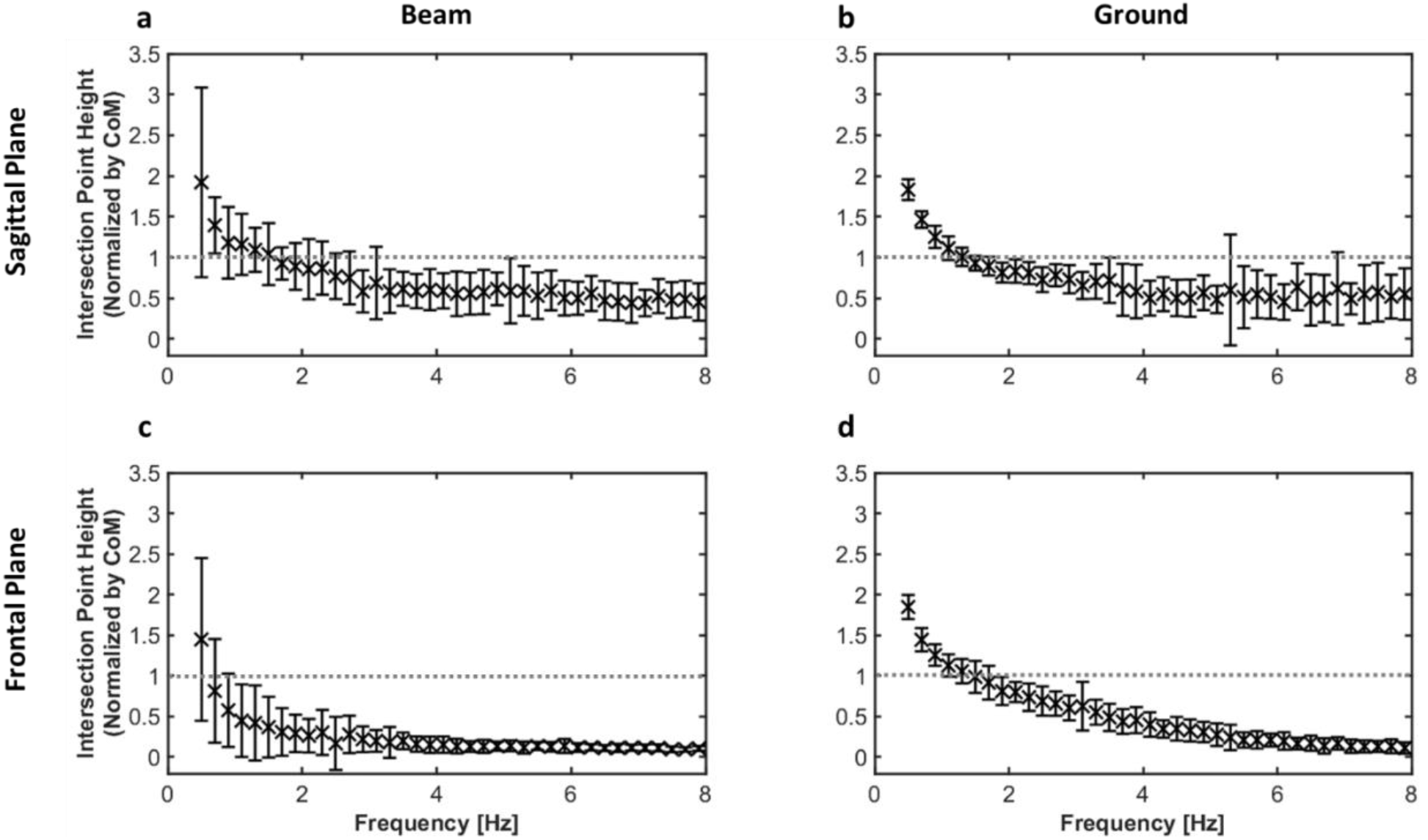
Human subject data of mean intersection point heights normalized by the center of mass height. The error bars for all plots indicate one standard deviation of the data. The center of mass height was computed for each subject. The normalized height of the center of mass is marked by a dashed line. **a)** Sagittal plane data of subjects standing in tandem stance on the beam. **b)** Sagittal plane data of subjects standing in tandem stance on the ground. **c)** Frontal plane data of subjects standing in tandem stance on the beam. **d)** Frontal plane data of subjects standing in tandem stance on the ground.

In contrast, differences in the intersection point’s frequency-dependent pattern arose in the frontal plane when standing on the beam and on the ground were compared. The intersection point was as high as the center of mass at 0.7 Hz on the beam and at 1.5 Hz on the ground. Additionally, at the highest analyzed frequency (8 Hz), the mean intersection point height was 0.11 on the beam and 0.14 on the ground. These results are depicted in Figure 6c and 6d. There are also visible differences in the variability across the conditions, where standing on the ground showed less interindividual variability than standing on a beam.

### Simulation Results

To analyze if unimpaired balance is best described by a controller that minimized effort, two values of *α*, very large (10^6^) for minimal control or very small (10^−4^) for “cheap” control, were compared. The controller that minimized joint torque effort with *α* **=** 10^6^ produced the best fit for the human results in both balance conditions and in both planes of analysis (Table 2). The best-fit controller reproduced all the human data, as evident from the overlap with the human data in Figure 7. These fits yielded root-mean-squared errors (RMSE) that are summarized in Table 2. In contrast, for *α* **=** 10^−4^, the smallest RMSE value was 0.088 for the ground condition in the frontal plane, which was 30% larger than the minimal control case for the same condition. Figure 8 shows that the error was overall larger when *α* was small, even though the *β* and *σ* _*r*_ parameters were selected to produce the smallest RMSE for each *α* condition.

**Table 2.** Best-fit parameters, simulated torque results, and RMSE.

| Platform | Plane | $\alpha$ | $\beta$ | $\sigma_r$ | RMS Torque Ratio | RMSE |
| --- | --- | --- | --- | --- | --- | --- |
|  |  | Large: min. ctrl. | <1: more hip penalty | <1: more hip noise | <1: more hip torque |  |
| Beam | Sagittal | $10^6$ | 0.2 | 1.0 | $3.30 \pm 0.08$ | 0.068 |
| Beam | Frontal | $10^6$ | 1.1 | 0.4 | $0.32 \pm 0.02$ | 0.070 |
| Ground | Sagittal | $10^6$ | 0.3 | 1.5 | $1.82 \pm 0.11$ | 0.072 |
| Ground | Frontal | $10^6$ | 0.2 | 0.01 | $2.74 \pm 0.02$ | 0.068 |

Values for RMS torque ratio are reported as mean ± SD.

**Figure 7.**
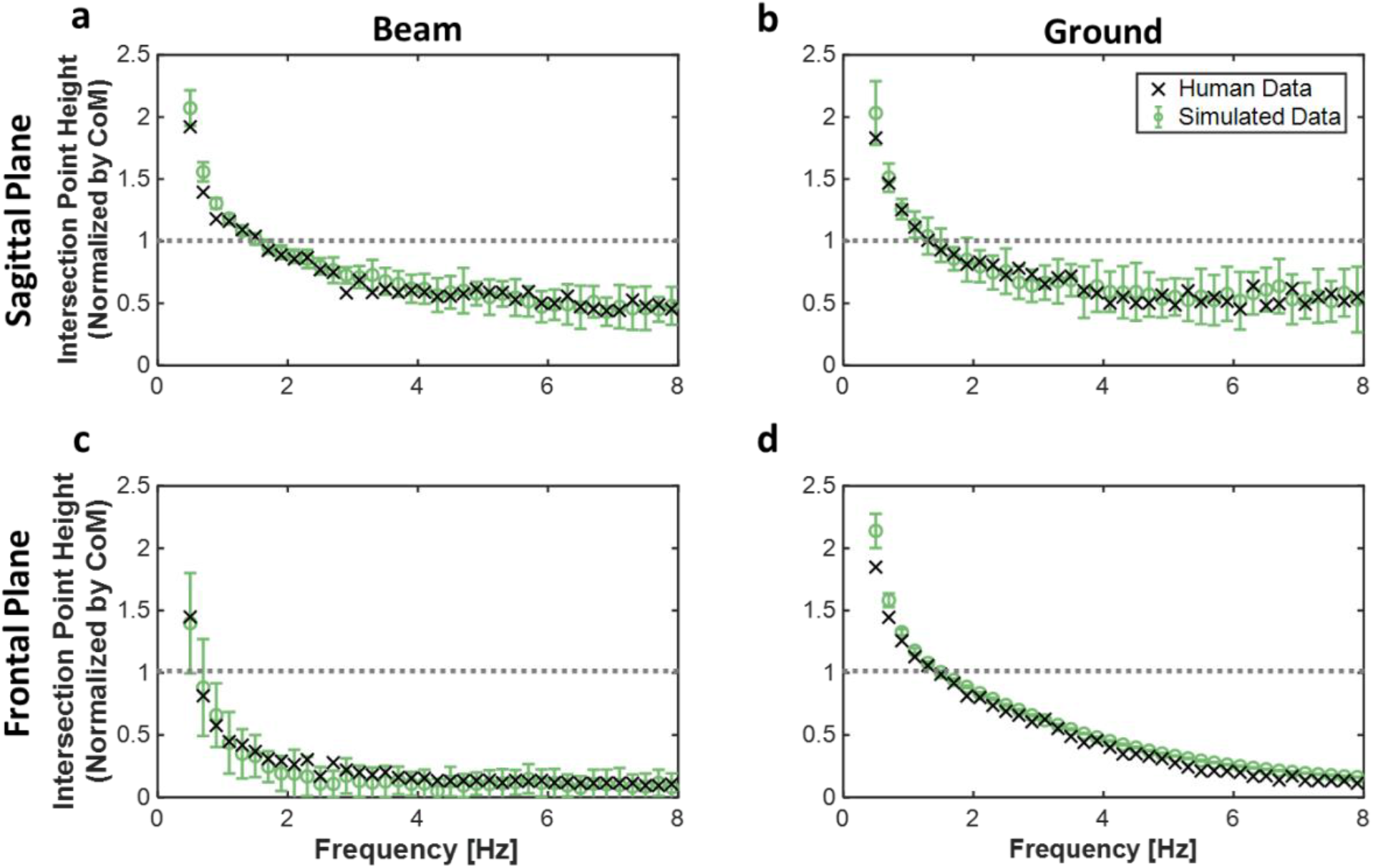
Mean intersection point heights normalized by the center of mass height. Human data are shown with symbol x, and the mean of the best-fit simulation results are shown with symbol o. The error bars denote the standard deviations across 40 simulation runs. The height of the center of mass is marked by a dashed line. For the human data, the center of mass height was computed for each subject; meanwhile, for the simulated data, the center of mass height was calculated based on the average height of the subjects and kept constant. **a)** Sagittal plane data of subjects standing in tandem stance on the beam. **b)** Sagittal plane data of subjects standing in tandem stance on the ground. **c)** Frontal plane data of subjects standing in tandem stance on the beam. **d)** Frontal plane data of subjects standing in tandem stance on the ground.

**Figure 8.**
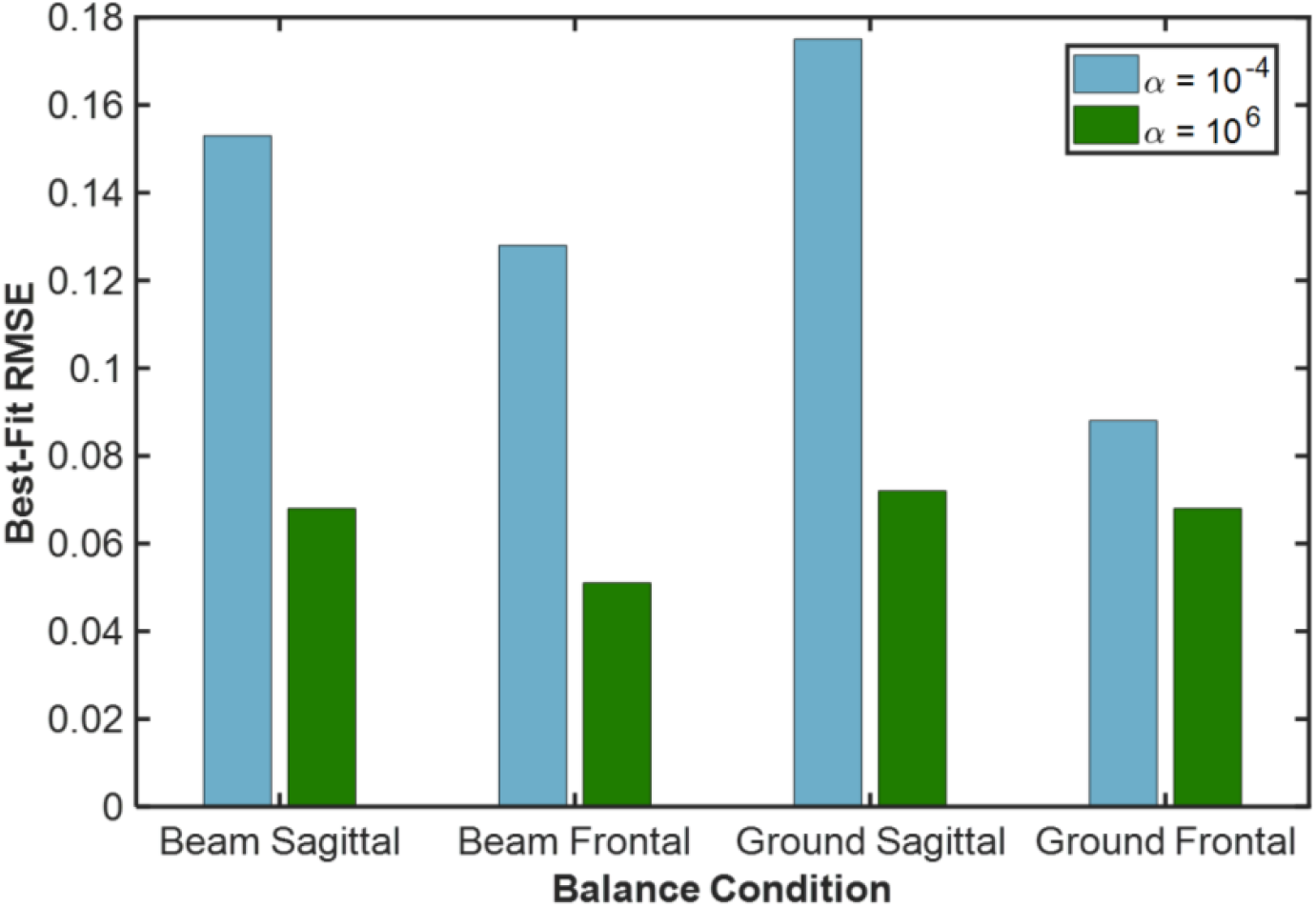
Best-fit RMSE values for small *α* (10^−4^) and large *α* (10^6^) in blue and green respectively, for each balance condition and for both planes. The best-fit RMSE was found by sweeping values of *β* and *σ* _*r*_, while setting *α* to be either large or small, then selecting the simulation result with the smallest RMSE.

Depending on the balance condition, the best-fit controllers varied in the way that they penalized the relative torque (*β*) and added noise to the ankle and hip joints (*σ* _*r*_). As shown in Figure 9, there was a range of *β* and *σ* _*r*_ combinations that produced small RMSEs for each condition and plane; the parameter sets that produced a particularly good fit are summarized in Table 2. Comparing the two standing conditions in the sagittal plane, the best-fit relative torque penalties were similar, and hip torque was penalized more than ankle torque. The root-mean-squared (RMS) torque ratio of greater than 1.00 agreed with this outcome and indicated that ankle torque was used more than the hip torque. The noise ratio *σ* _*r*_ was greater than or equal to 1.0 for both standing conditions in the sagittal plane.

**Figure 9.**
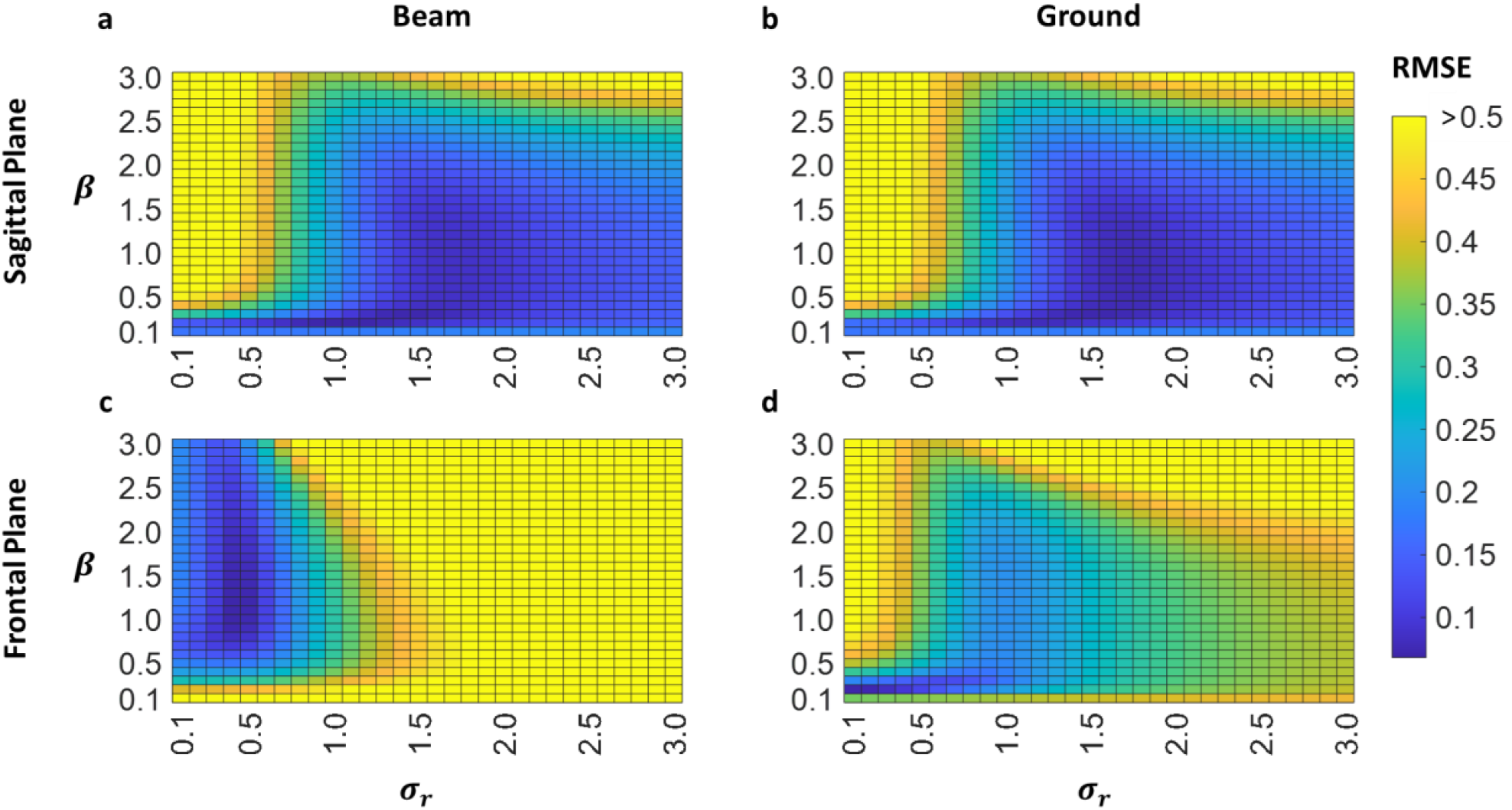
RMSE across a range of varied parameters for different balance conditions: **a)** sagittal plane data of subjects standing in tandem stance on the beam, **b)** sagittal plane data of subjects standing in tandem stance on the ground, **c)** frontal plane data of subjects standing in tandem stance on the beam, **d)** frontal plane data of subjects standing in tandem stance on the ground. The color bar indicates the RMSE values for all subplots. All RMSE values greater than 0.5 are represented by the same shade (yellow). *α* was set to be 10^6^. Some tested *β* and *σ* _*r*_ values were not reported in this graphic to maintain consistent intervals between each grid.

On the other hand, both the relative torque penalty, *β*, and the noise ratio, *σr*, differed between support conditions in the frontal plane. On the beam, the torque penalty ratio was greater than 1.0, suggesting that the ankle was more penalized than the hip. Agreeing with this outcome, the RMS torque ratio was less than 1.00 at 0.40, indicating that the hip was used more than the ankle when subjects stood on the beam. Meanwhile, on the ground, there was more hip penalty and use of ankle torque. The best-fit controller yielded more hip noise for both ground and beam conditions. On the ground, however, the weight of the noise on the hip was larger, with *σ* _*r*_ **=** 0.01. Finally, these variations in joint torque across conditions and planes of analyses consistently yielded best-fit of the human data with minimal control effort.

## DISCUSSION

This study set out to investigate balance control strategies of healthy young adults by using a challenging condition, standing on a narrow beam, as a probe. Simulations based on experimental data tested the hypothesis that subjects would minimize control effort while adjusting the relative joint effort penalty at the ankle and the hip. To this end, the intersection points of ground reaction forces at different frequency bands were extracted from the experimental data. While the overall trend of the intersection point’s frequency-dependent pattern was similar when standing on the beam versus on the ground in both planes of analysis, one notable variation was that the intersection point in the frontal plane with narrow support exhibited a different pattern. This result is consistent with the lateral instability evoked by standing on a narrow beam. Moreover, simulations of a simple nonlinear human balance model with a linear-quadratic optimal controller provided further insights from this experimental data. According to those results, the use of the ankle torque was penalized more than hip torque only on the beam in the frontal plane, which is likely a result of the mechanical constraints of the task as discussed below. The model results also showed that a controller that minimized overall joint torque effort closely replicated the human data for both balance conditions and in both planes of analysis.

### Human Results

The ground reaction force data of subjects standing in tandem stance on the beam and on the ground were analyzed in both the sagittal and frontal planes. Comparing the intersection point height patterns between conditions and planes of analysis, a common trend emerged: the intersection points were above the center of mass at low frequencies and below the center of mass at high frequencies. This pattern resembled the one previously shown in natural stance (parallel feet shoulder width apart) (18). Assuming that a balance controller is approximated by a double-inverted pendulum, at very low frequencies the body behaves like a single rigid body controlled at the ankle. Such behavior would correspond to an intersection point above the center of mass. If the double-inverted pendulum was stabilized solely by hip torque, the intersection point height would be zero. Both of these modes of motion have been reported in human data. Moreover, because this frequency-dependent pattern is the same regardless of foot placing, base of support, or plane of analysis, it likely reflects the basic biomechanical constraints of human balance (19).

To compare the intersection point profiles that emerged from balance on the beam and on the ground, it is useful to compare the planes of analyses separately. In the sagittal plane, regardless of the condition, both intersection point results exhibited a similar dependence on frequency. These findings are in line with previous work that investigated quiet, natural stance on the ground (18). On the other hand, this result is contrary to previous studies that reported different balance strategies (25) or increased postural sway (26) in the sagittal plane for different stance conditions. It was found that the ankle strategy dominated natural stance, while the hip strategy played a larger role in tandem stance (25); furthermore, tandem stance induced larger postural sway in the anteroposterior direction compared to natural stance (26). However, these previous results were based on measurements of the center of pressure alone. In tandem stance, it has been documented that individuals exhibited asymmetrical weight bearing, with the majority of the weight in the rear leg (27, 28). The discrepancy between our results and previous claims might arise from our analysis of force directions in addition to the center of pressure via the intersection point. Perhaps the observed difference in the center of pressure between natural and tandem stances is counteracted by distinct force directions in the two stances, resulting in similar intersection point patterns.

In contrast, in the frontal plane the high frequency asymptote of the intersection point curve shifted below that of the sagittal plane when subjects stood on the ground. As the intersection point gets closer to the ground, the body relies more on the hip for stabilization. Thus, the frontal plane data suggested that the hip’s role dominated at high frequencies compared to the sagittal plane case. More specifically, it is possible that the hip’s impedance played a large role, because higher frequency signals are correlated with interactive dynamics, rather than forward-path muscle dynamics (29).

Similarly in the frontal plane but when standing on a beam, the height of the high frequency asymptote also shifted closer to zero, and the frequency at which the intersection point height crossed the center of mass height was at a lower value. This finding suggested that the anti-phase mode of balance (or the “hip strategy”) was employed for most frequencies in the frontal plane. Previous findings (19) indicated that these subtle changes in the frequency-dependent intersection point suggest distinct controllers in the two balance conditions. To enhance insight, we employed a simple model with a stabilizing controller that was parameterized to best match the results to human data.

### Interpreting Best-Fit Controllers

To better understand how human control strategies differed or remained constant across standing conditions, we obtained controller parameters that both stabilized a double-inverted pendulum model and best described each balance condition’s frequency-dependent intersection point pattern. Note that this model could be fully controlled by ankle torque alone, by hip torque alone, or by both together (30). Therefore, an infinite number of ways existed to stabilize this system, yet the results suggested that only some joint torque combinations described the experimental data.

### Principle of Minimal Effort

Prior work on balance control suggested that unimpaired subjects minimized their overall control effort (3, 31, 32). To examine this prediction, the control parameter that determined the relative cost between state deviation and effort, α, was set to be either large (10^6^) or small (10^−4^), and the errors between the simulated results and human data were compared. Across both support conditions and in both planes of analysis, economized control, i.e., high α values, achieved the best fit. When *α* was large, the simulation result was not significantly affected by the choice of the design parameter that penalized state deviation from the nominal position. This feature greatly reduced the parameter space and simplified the analysis. Having employed a similar model, our previous study (19) showed that minimization of control effort could also describe human balance in the sagittal plane when subjects stood naturally. These results emphasize the competency of our model to describe a variety of stance conditions under one principle: minimized effort.

### Relative Joint Effort

While minimized control effort (large *α*) guaranteed a well-defined behavior of the closed-loop poles (unstable poles reflected across the imaginary axis), the control parameter that determined the relative penalty on the ankle and hip joint torques, *β*, affected the closed-loop control gains and thus the system response. Comparisons between the two balance conditions and the two planes of analyses suggested that the best-fit value of *β* varied in a condition-dependent manner.

In the sagittal plane, balancing in tandem stance was not significantly more difficult than balancing with parallel feet shoulder width apart because the base of support in the anteroposterior direction increased two-fold compared to placing one’s feet next to one another. Consistent with that account, the hip was similarly penalized more than the ankle for both tandem and natural stance conditions (19). On the other hand, the root-mean-squared torque ratio indicated that the role of the ankle was more dominant in tandem stance than natural stance. Furthermore, there was more ankle noise in tandem stance compared to natural stance. Given the assumption that force variability scaled linearly with mean force production (33), the noise in the ankle was expected to dominate in tandem stance more than in natural stance.

In the frontal plane, the best-fit control parameters varied depending on the platform conditions (on the ground vs. on the beam). In line with previous work (25, 27), on the ground, *β* was similar to that of the sagittal plane conditions and emphasized the use of the ankle torque over hip torque. On the contrary, when subjects stood on the beam, hip torque was used more than ankle torque on the beam. Subjects’ reliance on the hip joint when tasked with balance on a beam has also been reported in previous studies (34, 35). Perhaps because the beam constrained the center of pressure and induced a pressure distribution at the feet that would have been more uncomfortable than on the ground, control shifted to a more hip-dominant strategy.

Because task conditions were relatively challenging for this study, subjects were permitted to use their arms to maintain balance. Most subjects held out their arms to the side. This arm position shifted the moment of inertia compared to the case from a previous study when subjects stood on the ground with arms at the sides of the body (18, 19). As this difference in moment of inertia could contribute to the relative use of the ankle and hip joints, we modeled the arms as rigidly attached bars at the shoulders in this study. Moreover, previous work on beam walking has reported that subjects adopted a hip-dominant strategy in the mediolateral direction when their arms were allowed to move freely (34).

Although some subjects moved their arms, the simplified model used in this study was able to capture the subjects’ reliance on the hip when standing on the beam. Analysis of the effect of shoulder joint motion on balance is deferred to future work. These results showed that not only was the intersection point analysis sensitive to changes in the subjects’ posture and balance condition (mechanics), but it also allowed for the interpretation of healthy subject data under one central hypothesis: economizing effort. Although there were variations in the frequency-dependent intersection point across the tested conditions, the deliberately simplified model we used was able to capture all key features of the data with the available tuning parameters.

### Applications

As suggested above, the intersection point analysis provided insight into the relative roles of the ankle and hip joint torques. While kinematic-based methods like motion capture may be able to directly measure the movement of the body’s segments, the intersection point analysis only requires a force plate (assuming the center of mass height is a proportion of one’s height). Furthermore, as demonstrated above, the intersection point analysis enables the quantification of control strategies of balance. These features combined facilitate implementation of our methods to promote quantitative evaluation methods in a clinical setting.

A previous study showed that fall risk in older subjects increased with mediolateral balance impairments (36). While center-of-pressure-based measures in the anteroposterior direction (sagittal plane) were shown to differentiate between older and younger subjects (37–39), evidence for age-related changes in postural-sway-based measurements in the mediolateral direction (frontal plane) have been much weaker. As suggested in this study, the intersection point can differ between the sagittal and frontal planes. Thus, applying the intersection point analyses to studies on age-related changes may reveal new insights, specifically on the important topic of estimating fall risk.

### Limitations

Inevitably, limitations arise from using a simplified model. For example, the model incorporated two degrees of freedom, which was the minimum number of freely moving joints required to reproduce the experimental data (19). Furthermore, though the biomechanical model used in this study was nonlinear, the controller design was based on linearization. Consequently, the best-fit parameters reported are only valid for small motions near the upright position that are commonly observed during quiet standing. However, more challenging conditions, such as balancing on an unstable surface or even without vision, may evoke behavior that goes beyond small motions and may not be sufficiently well represented by linearized controllers. Nevertheless, the experimental data were well described by a linear model.

Finally, the simulations conducted in this work assumed highly simplified neurophysiology. For instance, neural transmission delay or sensory noise were excluded from this model for simplicity. The ankle and hip torques summarized the action of multiple muscles without any details of their contributions. By doing so, we were able to apply our proposed method to various balance conditions of unimpaired subjects. Future and ongoing work will implement this model to understand human data of other populations, such as older adults or persons with neural impairments (40), to investigate control strategies underlying the decline of balance ability.

### Conclusion

Incorporating the direction of the foot-ground reaction force to the location of the center of pressure yields a novel measure for evaluating upright stance performance: the height of the intersection point of ground reaction forces. By analyzing this intersection point’s frequency-dependent behavior, this work investigated balance control strategies involved in standing on and off a narrow beam. The results suggested that unimpaired subjects’ data is best described by a controller that minimizes overall effort, while penalizing the ankle and hip torques depending on the condition or the plane of analysis. Using a linearized control model could represent a range of conditions and showed the best fit when minimum control effort was assumed. This study provided new and actionable evidence that this control principle might be general for balance control, paving the way for future work to explore underlying neurological deficits and differences in subjects with impairments.

## GLOSSARY

*z*_*IP*_: Intersection point height
*x*_*CoP*_: Center of pressure position
*CoM*: Center of mass
*F*: Ground reaction force vector
*F*_x_: Horizontal component of the ground reaction force vector
*F*_*z*_: Vertical component of the ground reaction force vector
*θ*_*F*_: Ground reaction force vector direction
*τ*: Joint torque control input
*σ*_*r*_: Parameter that determines the ratio between noise on the ankle and hip joints; when it is set to be large, there is more noise in the ankle
*LQR*: Linear quadratic regulator
*R*: Design matrix that determines the cost of control effort
*α*: Parameter that determines the relative cost of control effort with respect to the cost on state deviation; when it is set to be large, minimal control is achieved
*β*: Parameter that determines the relative cost of control effort in the ankle compared to the hip joint; when it is set to be large, the ankle is penalized more than the hip
*RMSE*: Root-mean-squared error

## DATA AVAILABILITY

Data will be made available upon reasonable request.

ACKNOWLEDGMENTS

We would like to thank Christian Moses for his help in collecting the data. Marta Russo is currently at the Institute for Cognitive Sciences and Technologies (ISTC) at the National Research Council (CNR) in Rome, Italy.

## AUTHOR CONTRIBUTIONS

D.S. and N.H. contributed to conceiving and designing the research, M.R. performed human subject experiments and processed human subject data, K.S. performed simulation experiments and processed simulation data, J.L. developed the control model and simulations. All authors contributed to interpreting the results of experiments. K.S. and M.R. prepared the figures, K.S. drafted the manuscript, all authors edited and revised the manuscript, and all authors approved the final version of the manuscript.

## REFERENCES

1. Kiemel T, Zhang Y, Jeka JJ. Identification of neural feedback for upright stance in humans: Stabilization rather than sway minimization. J Neurosci 31: 15144–15153, 2011. doi: 10.1523/JNEUROSCI.1013-11.2011.

2. Van Der Kooij H, Van Asseldonk E, Van Der Helm FCT. Comparison of different methods to identify and quantify balance control. J Neurosci Methods 145: 175–203, 2005. doi: 10.1016/J.JNEUMETH.2005.01.003.

3. Kuo AD. An Optimal Control Model for Analyzing Human Postural Balance. IEEE Trans Biomed Eng 42: 87–101, 1995. doi: 10.1109/10.362914.

4. Lee J, Zhang K, Hogan N. Identifying human postural dynamics and control from unperturbed balance. J Neuroeng Rehabil 18: 1–15, 2021. doi: 10.1186/s12984-021-00843-1.

5. Winter DA, Patla AE, Ishac M, Gage WH. Motor mechanisms of balance during quiet standing. J Electromyogr Kinesiol 13: 49–56, 2003. doi: 10.1016/S1050-6411(02)00085-8.

6. Collins JJ, De Luca CJ. Open-loop and closed-loop control of posture: A random-walk analysis of center-of-pressure trajectories. Exp Brain Res 95: 308–318, 1993. doi: 10.1007/BF00229788.

7. Creath R, Kiemel T, Horak F, Peterka R, Jeka J. A unified view of quiet and perturbed stance: simultaneous co-existing excitable modes. Neurosci Lett 377: 75–80, 2005. doi: 10.1016/j.neulet.2004.11.071.

8. Ting LH. Dimensional reduction in sensorimotor systems: A framework for understanding muscle coordination of posture. Prog Brain Res 165: 299, 2007. doi: 10.1016/S0079-6123(06)65019-X.

9. Peterka RJ. Sensorimotor integration in human postural control. J Neurophysiol 88: 1097–1118, 2002. doi: 10.1152/jn.2002.88.3.1097.

10. Russo M, Lee J, Hogan N, Sternad D. Mechanical effects of canes on standing posture: beyond perceptual information. J Neuroeng Rehabil 19: 1–13, 2022. doi: 10.1186/s12984-022-01067-7.

11. Riemann BL, Guskiewicz KM, Shields EW. Relationship between Clinical and Forceplate Measures of Postural Stability. J Sport Rehabil 8: 71–82, 1999. doi: 10.1123/JSR.8.2.71.

12. Promsri A, Haid T, Werner I, Federolf P. Leg Dominance Effects on Postural Control When Performing Challenging Balance Exercises. Brain Sci 2020, Vol 10, Page 128 10: 128, 2020. doi: 10.3390/BRAINSCI10030128.

13. Sozzi S, Honeine JL, Do MC, Schieppati M. Leg muscle activity during tandem stance and the control of body balance in the frontal plane. Clin Neurophysiol 124: 1175–1186, 2013. doi: 10.1016/J.CLINPH.2012.12.001.

14. Tanaka H, Uetake T, Kuriki S, Ikeda S. Changes in Center-of-Pressure Dynamics during Upright Standing Related to Decreased Balance Control in Young Adults: Fractional Brownian Motion Analysis. J Hum Ergol (Tokyo) 31: 1–11, 2002. doi: 10.11183/JHE1972.31.1.

15. Ersal T, Sienko KH. A mathematical model for incorporating biofeedback into human postural control. J Neuroeng Rehabil 10: 1–13, 2013. doi: 10.1186/1743-0003-10-14.

16. Duarte M, Freitas SMSF. Revision of posturography based on force plate for balance evaluation. Brazilian J Phys Ther 14: 183–192, 2010. doi: 10.1590/S1413-35552010000300003.

17. Duarte M, Sternad D. Complexity of human postural control in young and older adults during prolonged standing. Exp Brain Res 191: 265–276, 2008. doi: 10.1007/s00221-008-1521-7.

18. Boehm WL, Nichols KM, Gruben KG. Frequency-dependent contributions of sagittal-plane foot force to upright human standing. J Biomech 83: 305–309, 2019. doi: 10.1016/j.jbiomech.2018.11.039.

19. Shiozawa K, Lee J, Russo M, Sternad D, Hogan N. Frequency-dependent force direction elucidates neural control of balance. J Neuroeng Rehabil 18: 1–12, 2021. doi: 10.1186/s12984-021-00907-2.

20. O’Connor SM, Kuo AD. Direction-dependent control of balance during walking and standing. J Neurophysiol 102: 1411–1419, 2009. doi: 10.1152/jn.00131.2009.

21. Winter DA, Patla AE, Prince F, Ishac M, Gielo-perczak K. Stiffness control of balance in quiet standing. J Neurophysiol 80: 1211–1221, 1998. doi: 10.1152/jn.1998.80.3.1211.

22. Loram ID, Lakie M. Human balancing of an inverted pendulum: position control by small, ballistic-like, throw and catch movements. J Physiol 540: 1111–1124, 2002. doi: 10.1113/JPHYSIOL.2001.013077.

23. De Leva P. Adjustments to zatsiorsky-seluyanov’s segment inertia parameters. J Biomech 29: 1223–1230, 1996. doi: 10.1016/0021-9290(95)00178-6.

24. Sugimoto-Dimitrova R, Shiozawa K, Gruben K, Hogan N. Frequency-domain patterns in foot-force line-of-action: an emergent property of standing balance control. .

25. Winter DA, Prince F, Frank JS, Powell C, Zabjek KF. Unified theory regarding A/P and M/L balance in quiet stance. J Neurophysiol 75: 2334–2343, 1996. doi: 10.1152/JN.1996.75.6.2334.

26. Kenny RPW, Eaves DL, Martin D, Hatton AL, Dixon J. The effects of textured insoles on quiet standing balance in four stance types with and without vision. BMC Sports Sci Med Rehabil 11: 1–8, 2019. doi: 10.1186/s13102-019-0117-9.

27. Jonsson E, Seiger Å, Hirschfeld H. Postural steadiness and weight distribution during tandem stance in healthy young and elderly adults. Clin Biomech 20: 202–208, 2005. doi: 10.1016/J.CLINBIOMECH.2004.09.008.

28. Nichols DS, Glenn TM, Hutchinson KJ. Changes in the mean center of balance during balance testing in young adults. Phys Ther 75: 699–706, 1995. doi: 10.1093/PTJ/75.8.699.

29. Zahalak GI, Heyman SJ. A Quantitative Evaluation of the Frequency-Response Characteristics of Active Human Skeletal Muscle In Vivo. J Biomech Eng 101: 28–37, 1979. doi: 10.1115/1.3426220.

30. Murray R, Hauser J. A Case Study in Approximate Linearization: The Acrobot Example. .

31. Kiemel T, Zhang Y, Jeka JJ. Identification of neural feedback for upright stance in humans: Stabilization rather than sway minimization. J Neurosci 31: 15144–15153, 2011. doi: 10.1523/JNEUROSCI.1013-11.2011.

32. Versteeg CS, Ting LH, Allen JL. Hip and ankle responses for reactive balance emerge from varying priorities to reduce effort and kinematic excursion: A simulation study. J Biomech 49: 3230–3237, 2016. doi: 10.1016/J.JBIOMECH.2016.08.007.

33. Jones KE, Hamilton AF d. C, Wolpert DM. Sources of signal-dependent noise during isometric force production. J Neurophysiol 88: 1533–1544, 2002. doi: 10.1152/jn.2002.88.3.1533.

34. Huber ME, Chiovetto E, Giese M, Sternad D. Rigid soles improve balance in beam walking, but improvements do not persist with bare feet. Sci Rep 10, 2020. doi: 10.1038/S41598-020-64035-Y.

35. Otten E. Balancing on a narrow ridge: biomechanics and control. Philos Trans R Soc B Biol Sci 354: 869, 1999. doi: 10.1098/RSTB.1999.0439.

36. Maki BE, Holliday PJ, Topper AK. A Prospective Study of Postural Balance and Risk of Falling in An Ambulatory and Independent Elderly Population. J Gerontol 49: M72–M84, 1994. doi: 10.1093/GERONJ/49.2.M72.

37. Maki BE, Holliday PJ, Fernie GR. Aging and postural control. A comparison of spontaneous- and induced-sway balance tests. J Am Geriatr Soc 38: 1–9, 1990. doi: 10.1111/J.1532-5415.1990.TB01588.X.

38. Prieto TE, Myklebust JB, Hoffmann RG, Lovett EG, Myklebust BM. Measures of postural steadiness: differences between healthy young and elderly adults. IEEE Trans Biomed Eng 43: 956–966, 1996. doi: 10.1109/10.532130.

39. Lizama LEC, Pijnappels M, Faber GH, Reeves PN, Verschueren SM, Van Dieën JH. Age Effects on Mediolateral Balance Control. PLoS One 9: e110757, 2014. doi: 10.1371/JOURNAL.PONE.0110757.

40. Bartloff JN, Ochs WL, Nichols KM, Gruben KG. Frequency-dependent behavior of paretic and non-paretic leg force during standing post stroke. .

